# Reader Engagement with Wikipedia’s Medical Content

**DOI:** 10.1101/797779

**Authors:** Lauren A. Maggio, Ryan M. Steinberg, Tiziano Piccardi, John M. Willinsky

## Abstract

Wikipedia’s extensive health and medical entries, maintained by WikiProject Medicine (WPM), are well supported by external links that provide readers with both a means of verifying the sources drawn upon and visiting those sources to learn more about a topic. In analysing how readers approach these links, data was collected on reader engagement with these links on WPM pages and on the rest of Wikipedia over a 32-day period. Readers of WPM pages were found to engage with external links more frequently than readers of the rest of Wikipedia, with WPM readers favoring hovering over a link and footnote clicking, compared to W readers who tended to click more external links per page viewed. Compared to readers of the rest of Wikipedia, WPM readers appear more attentive to the external link’s function in verifying and authorizing Wikipedia content, than to the educational potential of examining the sources themselves.

## Introduction

Wikipedia is a freely available online encyclopedia that intends to provide “every single person on the planet free access to the sum of all human knowledge” (Wikimedia Foundation, 2004). To meet this mission, thousands of volunteer editors have created almost six million English-language Wikipedia pages many of which include hyperlinked footnotes to the sources they draw upon in assembling and verifying the content (<u>Wikipedia:Statistics</u>, 2019). These linked references not only serve as sources for Wikipedia content, lending it authority (Fallis 2008), but offer readers a gateway to further learning, with this opportunity enhanced by the growing degree of public access to research literature (Piwowar et al., 2018).

Wikipedia is proving to be a leading source of health information (Heilman et al., 2015; Laurent et al., 2009), and reader engagement with a page’s references can provide opportunities to understand a diagnosis or inform a conversation with their physician. Wikipedia’s health-focused pages, which are maintained by <u>WikiProject Medicine</u> (WPM) editors, are thought to meet a high standard for quality and rigor (James, 2016; Maskalyk, 2014; Trevena, 2011). For example, WPM recommends that editors select “reliable sources” for the content they present, with a favoring of “review articles (especially systematic reviews) published in reputable medical journals” (<u>Wikipedia:Identifying and using primary sources</u>, 2019). This is particularly important for health professionals and students, which research shows are active Wikipedia readers (Scafidi et al., 2019; Egle et al., 2015; Allawahla et al., 2013). They are familiar with the research literature, and expected to engage in evidence-based medicine, which involves analyzing and applying evidence (e.g., research from systematic reviews, randomized trials) in patient care (Guyatt et al., 1992). To this end, faculty at several health professions schools teach courses on editing Wikipedia (Joshi et al., 2019; Appollonio et al., 2018).

In a previous study, we investigated Wikipedia’s gateway effect by tracking referrals from Wikipedia to research article DOIs (Digital Object Identifier) through Crossref, combined with page view information from Wikimedia (Maggio et al., 2017). However, with the data available at the time, we could not determine if a click to an external reference with a DOI originated from a WPM page.

Thus, to learn more about reader engagement with Wikipedia’s external references, this study leverages Wikimedia’s newly created infrastructure for data collection to compare engagement with external links by readers of WPM pages with that of readers of the rest of Wikipedia. The study is guided by two research questions:

RQ 1: How does reader engagement with references in WPM articles compare to engagement in the rest of Wikipedia?

RQ 2: To what extent do readers’ patterns of engagement speak to their regard for WPM’s sources as validating Wikipedia content and/or as a gateway to further learning?

## Method

With the approval and support of the Wikimedia Foundation (WMF), we collected the data presented in this study between March 22 - April 22, 2019 from <u>Wikimedia’s Event Logging</u> system and the production MediaWiki database, with the data remaining within that system, as required by WMF, for a year before being deleted (see Acknowledgements). We anonymized the aggregated data by removing IP addresses, identifying browser information, and reader-sessions associated with page edits. While this data is not publicly available, researchers may request access by submitting a request to the <u>WMF research team</u>. The code utilized to collect and analyze the data, however, is organized and made publicly available in a collated series of Jupyter notebooks at: https://github.com/ryanmax/wiki-citation-usage/blob/master/README.md (Steinberg et al., 2019).

### Data Collection

The data is drawn from English Wikipedia pages in the <u>main namespace</u>, a designation that contains the encyclopedia proper. Wikipedia pages or topics were identified as being part of two main groupings: WikiProject Medicine pages (WPM) and the rest of Wikipedia (W). To be included in the study, the pages from each grouping had to have at least one external link in the <u>externallinks</u> table. The <u>categorylinks</u> table was used to define the WPM pages, with each possessing a Talk page bearing the category “All WikiProject Medicine articles.” Both the externallinks and categorylinks tables were queried twice (April 1, 2019 and April 20, 2019) during 32-day study period (March 22 - April 22, 2019).

For determining the number of pages, length of pages, the number of external links, and the number of “freely accessible” links that editors have added as the sources of their work, a single day’s worth of database and XML dump files was captured from late in the study period (April 20, 2019), and which, as it had only 0.5 percent more external links than on April 1, 2019, was felt to be sufficiently representative to serve as the source for all static data counts. The external link count, which is based on MediaWiki’s <u>externallinks</u> table, does not include <u>interwiki links</u>, representing abbreviated forms of commonly-used internal and external links, which limits the accuracy of external link counts for both WPM and W. The event logging system this study relied on similarly omitted data from interwiki links, meaning the definition of an external link used across this study is consistent.

Page view data was gathered from the <u>wmf.pageview_hourly</u> table. WMF employs methods to identify bot traffic in page view data, which was excluded in our analysis. In reporting the data collected over a 32-day period (March 22 - April 22, 2019), the raw counts were divided by 32 to create a count approximating a “daily average” for these counts, in light of this serving as a common measure of internet traffic.

Reader engagement with external links was gathered from <u>Wikimedia’s Event Logging system</u> using the <u>CitationUsage schema</u>, instrumented by Wikimedia’s programmers, following a month of piloting and refinement for this study. The CitationUsage schema collected all sessions with reader engagement, except for those involving anonymous Wikipedia editors (21 sessions out of 72,953,065 total which translated into the removal of 34 citation events out of a total of 113,520,376). For the entire study period (March 22 - April 22, 2019), the CitationUsage schema captured the following reader “events”: (a) clicking an external link; (b) clicking on a reference link listed among a page’s set of references; (c) clicking a footnote link leading to a reference on the page; (d) hovering over a reference link; and time from page view to event (Table 1). Additionally, the CitationUsage schema captured the clicking of links bearing the “freely accessible” icon.

**Table 1.**
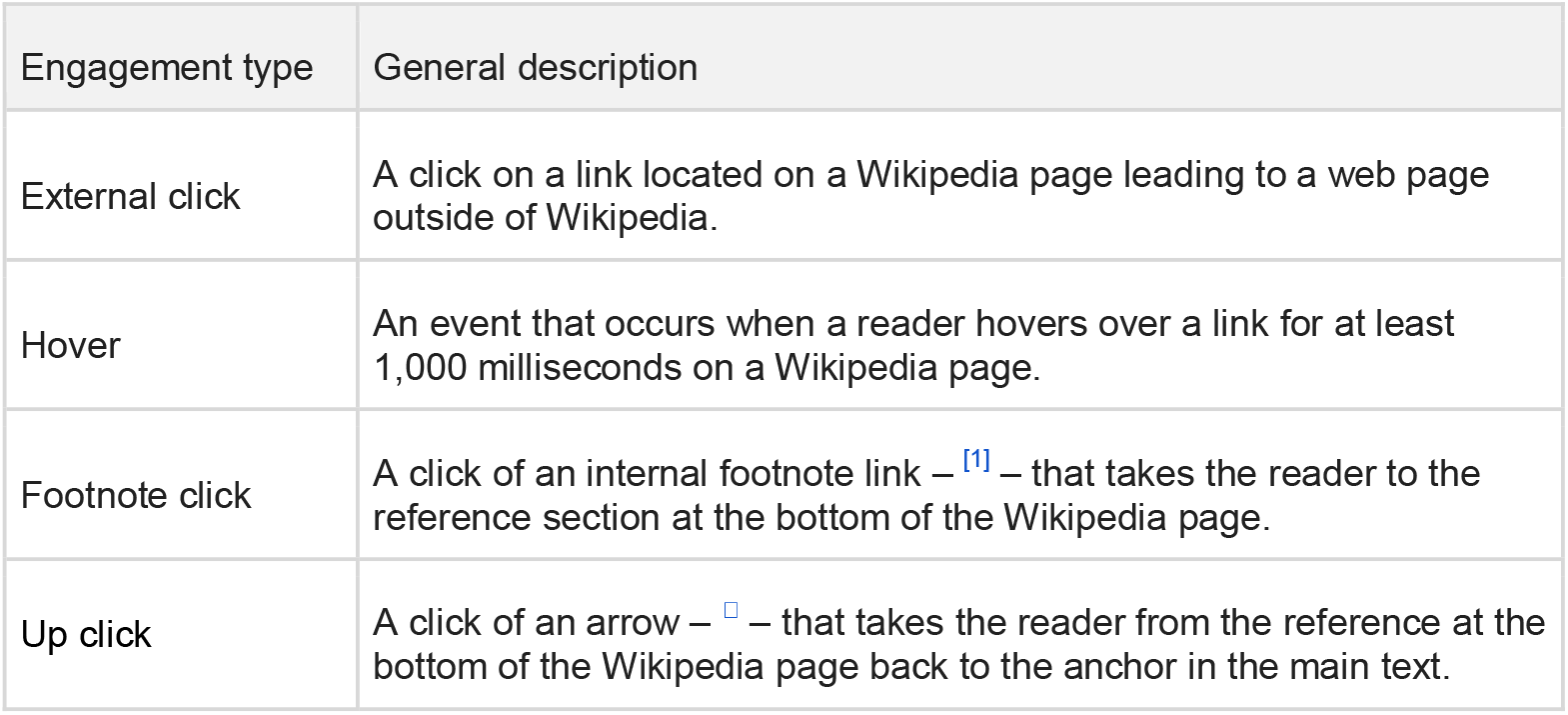
Types of link events captured by the CitationUsage Schema.

| Engagement type | General description |
| --- | --- |
| External click | A click on a link located on a Wikipedia page leading to a web page outside of Wikipedia. |
| Hover | An event that occurs when a reader hovers over a link for at least 1,000 milliseconds on a Wikipedia page. |
| Footnote click | A click of an internal footnote link – <sup>[1]</sup> – that takes the reader to the reference section at the bottom of the Wikipedia page. |
| Up click | A click of an arrow – – that takes the reader from the reference at the bottom of the Wikipedia page back to the anchor in the main text. |

### Analysis

Descriptive statistics were calculated using Excel (Redmond, WA). Inferential statistics were calculated to determine the significance in the difference between engagements in WPM and W using the Python library SciPy. In our analysis, we verified the statistical differences between (a) WPM and W in the <u>time to first engagement</u> with a link, (b) in the <u>length of the articles</u>, (c) and in the number of <u>page loads per event</u>. In the first two cases (a) (b), we used a normality test to determine the type of statistical test to apply. In both comparisons, after rejecting the normality hypothesis, we applied Mann-Whitney U-Test, a non-parametric test that does not require normality assumptions on the two distributions. In the third case (c), we compared the two groups over all possible combinations of access method and type of event with a two-tailed Fisher’s exact test. To explore and visualize the data, we used respectively Spark/Pandas and the library Seaborn. The complete analysis is available in the project’s publicly accessible Github repository (Steinberg et al., 2019).

### Stakeholder engagement

This study was possible through collaboration with the WMF research team beginning in 2018, with our <u>publishing of a research proposal</u> to Wikimedia’s Meta-Wiki (Maggio et al., 2019). Members of the Wikipedia community, including members of WPM, were invited to provide feedback and pose questions. Over several months, we held teleconferences and in-person meetings with the WMF research team to understand Wikimedia’s infrastructure, parameters of data use (e.g., best practices for accessing the data securely; duration of data access), and the expectations of the Wikipedia community. Prior to data collection, a WMF research team member posted to Wikipedia’s Village Pump, a set of Wikipedia pages dedicated to discussing technical issues and policies, an <u>announcement</u> of the data collection (Redi, 2019). The announcement invited the Wikipedia community to post public comments and provided contact information for expressing concerns about the research.

## Results

### Wikipedia Pages and External Links

This study compares readers’ engagement with the pages curated by WikiProject Medicine (WPM) to their engagement with the rest of the English edition of Wikipedia (W). At the time of this study, WPM represented 34,324 pages (i.e., subject or topic entries), while the rest of W had 5,839,083 pages (Table 2). WPM pages possessed more than twice as many links to external references and other sites than the rest of Wikipedia. WPM pages were 13,084.9 characters in length on average (SD=19,378.4; median=6,628.0; IQR=11,640), which was 70.1 percent longer than the average page in the rest of Wikipedia at 7,676.4 characters (SD=13,632.4; median=3,865.0; IQR=5,789) (Fig. 1). WPM pages also demonstrated a greater “link density” with an external link appearing every 450.3 characters per page, compared to a link every 657.3 characters for the rest of Wikipedia.

**Table 2.** Wikipedia pages, page length, and external links for WikiProject Medicine pages (WPM) and the rest of Wikipedia (W) on April 20th, 2019.

|  | WPM | W |
| --- | --- | --- |
| Wikipedia pages | 34,324 | 5,839,083 |
| Pages with external links | 32,609 | 5,210,746 |
| External links | 945,645 | 60,851,396 |
| Links per page (with links) | 29.0 | 11.7 |
| Page length (characters) | 13,084.9 | 7,676.4 |
| Characters per link for pages (with links) | 450.3 | 657.3 |

**Figure 1.**
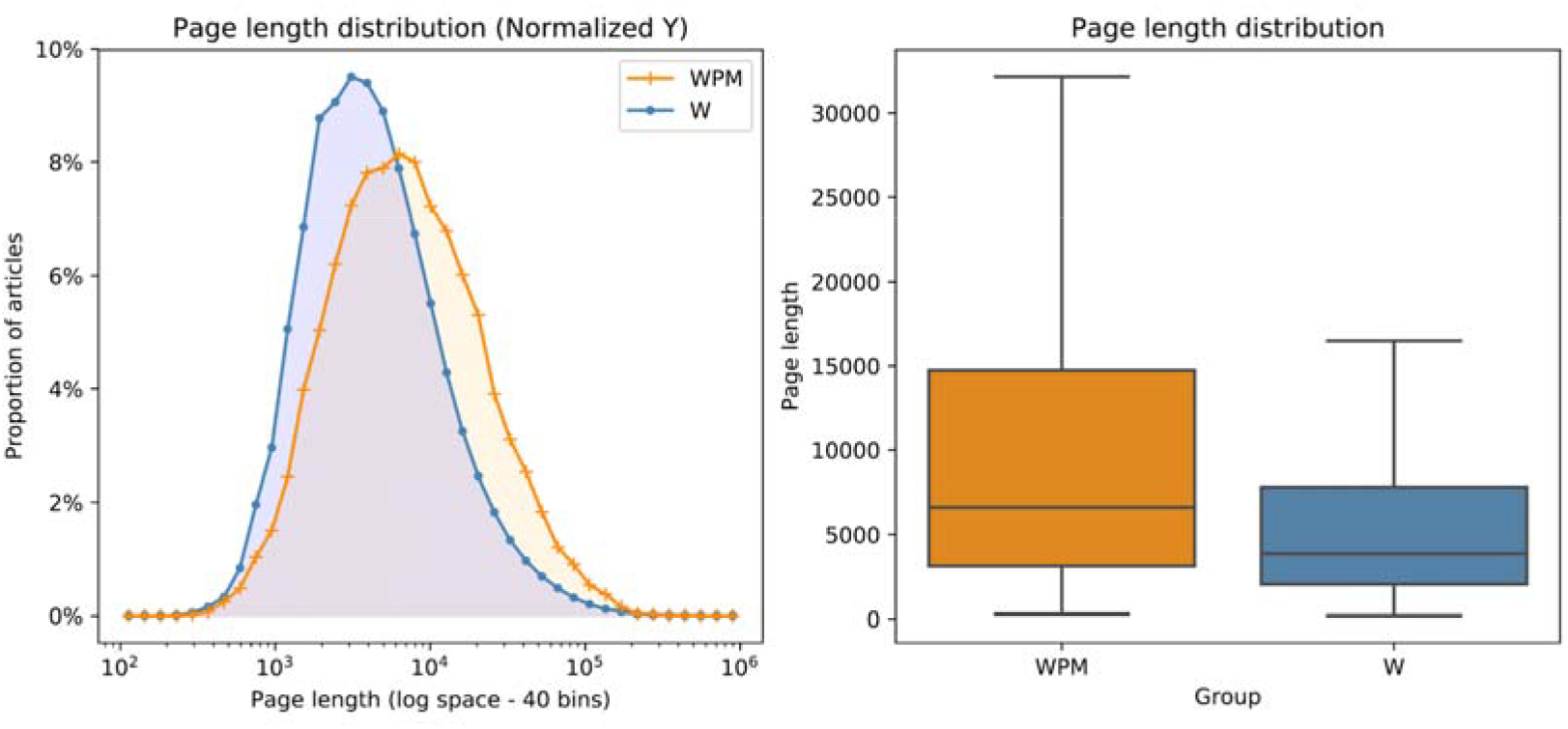
Distribution of page lengths in characters for WikiProject Medicine pages (WPM) and the rest of Wikipedia (W) on April 20th, 2019. The difference between the two distributions is statistically significant according to Mann–Whitney U test (P << 0.001, two-tailed).

The biomedical literature, including research reviews and studies, was a leading source of WPM external links. For example, the most common hostname was “www.ncbi.nlm.nih.gov,” accounting for 25.2 percent of the WPM’s external links, indicating the likely citing of a research paper found through the PubMed database run by the National Institutes of Health (NIH). It was followed by the hostnames “www.worldcat.org” with 2.7 percent and “www.google.com” with 1.2 percent of the external links. As for the hostnames that prevailed among external links in the rest of Wikipedia, the three leading hostnames were “tools.wmflabs.org” (where Wikimedia hosts <u>tools</u> developed to assist editors) at 3.5 percent, “www.google.com” at 2.9 percent and “books.google.com” at 2.0 percent, while “www.ncbi.nlm.nih.gov” still accounted for 1.8 percent of the external links outside of WPM.

### Page Views and Events

Readers viewed 5,875,470.4 WPM pages a day on average, compared to 228,445,128.4 pages for readers of the rest of Wikipedia (Table 3). While WPM’s page count is 0.6 percent of W’s count, the number of page views that WPM received from readers was 2.6 percent of that for W. This suggests that WPM pages were viewed more than four times as frequently as the rest of Wikipedia. The majority of these views for both WPM and W took place on mobile devices, with WPM readers the heavier users of mobile devices in accessing Wikipedia.

**Table 3:** Daily average of page views by device for WikiProject Medicine (WPM) pages and the rest of Wikipedia (W) for March 22 to April 22, 2019.

|  | WPM (%) | W (%) |
| --- | --- | --- |
| Page views on desktop device | 1,957,821.6 (33.3) | 97,956,273.1 (42.9) |
| Page views on mobile device | 3,917,648.8 (66.7) | 130,488,855.3 (57.1) |
| Page view total | 5,875,470.4 (100) | 228,445,128.4 (100) |

Readers spent more time on a page before first clicking an external link after loading a WPM page. The WPM reader’s median time of 47.5 seconds (SD=1.2; mean=82.5; IQR=147,306) was 44.8% longer than the median time of 32.8 seconds (SD=5.6; mean=39.1; IQR=86,424) for W readers. The difference between the two distributions is statistically significant according to Mann–Whitney U test (P << 0.001, two-tailed).

Among the four types of events recorded, WPM readers (Table 4) were more likely to, above all, hover over a footnote or other link, especially on their desktop devices, as well as more likely to click on footnote links, compared to W readers (Table 5). On the other hand, W readers were more likely to click on external links than WPM readers, favoring their mobile devices in that regard. The WPM readers are interested, it appears, in seeing what evidence is being drawn upon in making the statements set out on a WPM page, but not as given to clicking on such external links to explore what they have to offer as readers of the rest of Wikipedia.

**Table 4:** Daily average of event type (actions taken by readers) for WikiProject Medicine (WPM) by device for March 22 to April 22, 2019.

| Event type | WPM total (%) | WPM desktop | WPM mobile (%) |
| --- | --- | --- | --- |
| Hover over link | 48,748.9 (46.9) | 45,814.8 (60.3) | 2,934.1 (10.5) |
| Footnote click | 27,739.4 (26.7) | 10,948.8 (14.4) | 16,790.6 (60.3) |
| External Click | 25,811.9 (24.9) | 17,792.3 (23.4) | 8,019.7 (28.8) |
| Up click | 1,539.5 (1.5) | 1,422.8 (1.9) | 116.4 (0.4) |
| All events | 103,839.7 (100) | 75,978.7 (100) | 27,860.8 (100) |

**Table 5:** Daily average of event type (actions taken by readers) for Wikipedia (W) by device for March 22 to April 22, 2019.

| Event type | W total (%) | W desktop (%) | W mobile (%) |
| --- | --- | --- | --- |
| Hover over link | 1,122,704.0 (32.7) | 1,057,982.0 (47.2) | 64,722.0 (5.4) |
| Footnote click | 722,131.0 (21.0) | 235,245.0 (10.5) | 486,886.7 (40.7) |
| External Click | 1,557,125.0 (45.3) | 915,445.1 (40.9) | 641,676.4 (53.6) |
| Up click | 34,738.0 (1.0) | 31,230.1 (1.4) | 3,508.1 (0.3) |
| All events | 3,436,698.0 (100) | 2,239,902.2 (100) | 1,196,793.2 (100) |

Among the external links in WPM, 22.1% of the research citations drawn from PubMed are labeled as “freely accessible”, and on desktop devices include a link bearing an open access icon (a green open lock, with the rollover label “freely accessible”), although no evidence was found that links with icons were clicked more frequently than without it (Fig. 2). This may be, in part, because the typical research citation has three or four links leading to (a) the article on the publisher’s site (doi link); (b) its PubMed entry (PMID link), and, if open access, the article in PubMed Central (PMC link and article title link).

**Figure 2.**
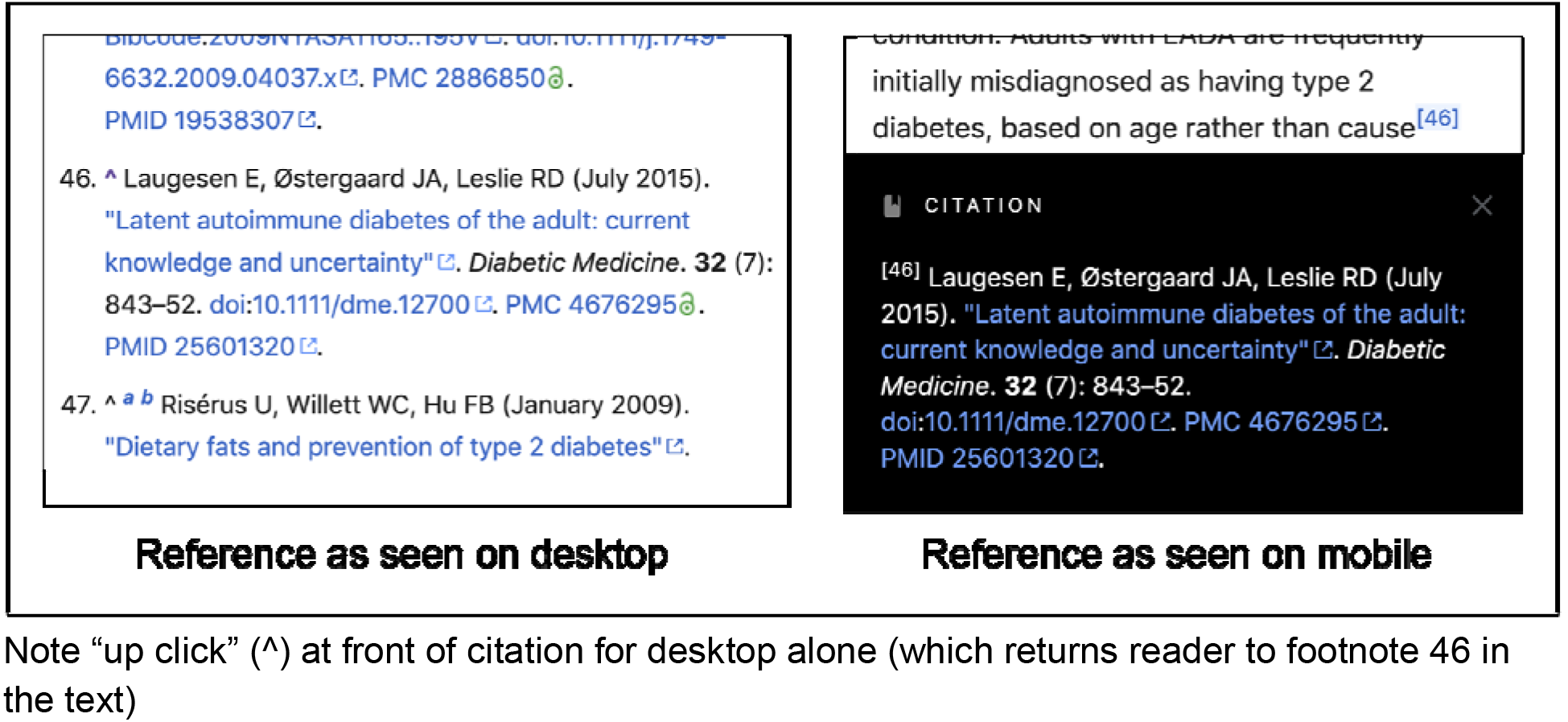
A research study cited on Wikipedia’s “diabetes” page displayed on desktop device with “freely accessible” icon for PubMed Central (PMC) link and on mobile device without open access icon.

To further compare event data between WPM and W, the number of page views per event was calculated to determine, in effect, how many pages readers viewed before engaging with an external link (Table 6). Overall, readers of WPM pages were more likely (56.6 page views per event) to engage with a link than were W readers (66.5 page views per event). The differences in engagement were most pronounced with WPM readers hovering over a link on their desktop device. Yet W readers more frequently clicked external links, with the difference especially notable when on their mobile device, where they were almost twice as likely to click an external link than WPM readers (suggesting that the external links in WPM are not mobile-friendly). The up click, in which a reader clicks the link from a footnote back (up) to the text, was not a function that appeared on the iOS and Android mobile devices we tested, as the footnote appears at the bottom of the screen when clicked and disappears on touching the text.

**Table 6:**
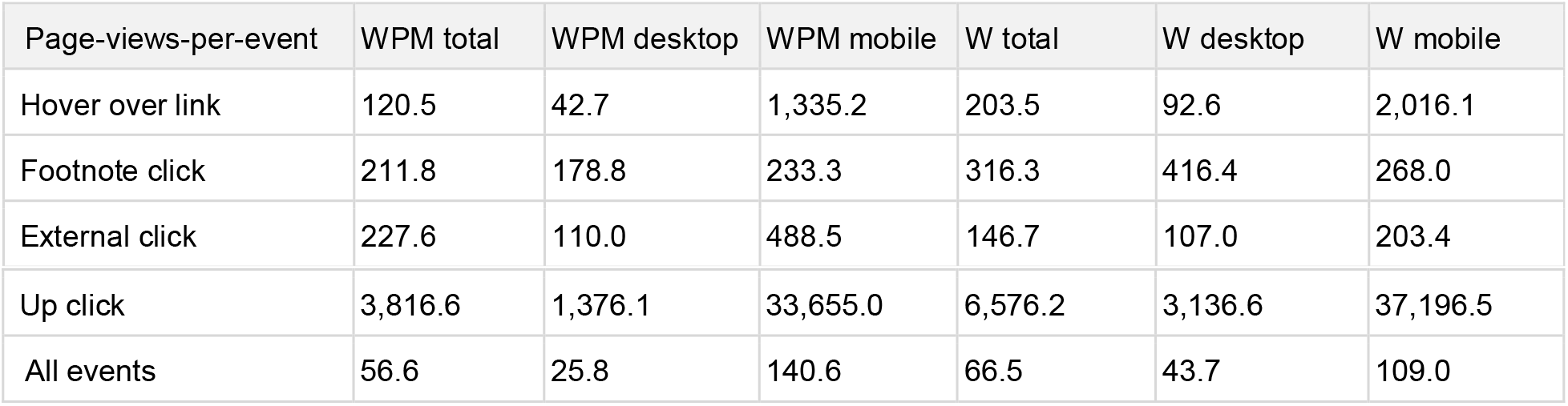
Page views per event for WikiProject Medicine (WPM) and Wikipedia (W) for March 22 to April 22, 2019. The difference between each pair of WPM and W distributions is statistically significant as derived from Fisher’s exact test (*p* < 0.001, two-tailed).

| Page-views-per-event | WPM total | WPM desktop | WPM mobile | W total | W desktop | W mobile |
| --- | --- | --- | --- | --- | --- | --- |
| Hover over link | 120.5 | 42.7 | 1,335.2 | 203.5 | 92.6 | 2,016.1 |
| Footnote click | 211.8 | 178.8 | 233.3 | 316.3 | 416.4 | 268.0 |
| External click | 227.6 | 110.0 | 488.5 | 146.7 | 107.0 | 203.4 |
| Up click | 3,816.6 | 1,376.1 | 33,655.0 | 6,576.2 | 3,136.6 | 37,196.5 |
| All events | 56.6 | 25.8 | 140.6 | 66.5 | 43.7 | 109.0 |

## Discussion

The net outcome of this analysis is that readers of WPM appear to be less likely to explore what can be learned from the external links they encounter in reading WPM pages than readers of the rest of Wikipedia, even as they are exposed to more of these external links in the course of their reading. The relatively higher frequency of WPM reader visits points to both the level of trust that WPM readers have in Wikipedia’s coverage of health and medical topics and their interest in continuing to bolster and validate this trust by viewing WPM citations.

These conclusions suggest that the WPM citation’s predominant function is a trust-building in, and validation of, the page’s content. This is a necessary aspect to offering an unconventional, relatively recently formed source of information, such as Wikipedia, in an area of such vital import. The plenitude of citations speaks to a transparency around the construction of authority that marks the Wikipedia project. Yet given the power of such references in a digital setting to not only reassure but educate readers, this study also speaks to what might be cast as missed opportunities for further learning and deeper inquiry into the topics being consulted, a possibility only heightened by the demonstrated use of WPM by medical students and practicing physicians (Scafidi et al., 2019; Rossler et al., 2015; Allawahla et al., 2013).

As the use of Wikipedia continues to grow, the additional levels of trust to be gained from hovering over references will suffer diminishing returns. This prospect should lead us to further considerations of what stands in the way of WPM pages serving as a greater gateway to further learning. Potential reasons for WPM readers being less likely to click on external links than readers of the rest of Wikipedia may include:

1. Readers may turn to WPM for quick answers in pressing situations (e.g., to check for signs and symptoms of an emergent condition), and not have time to, or interest in, clicking on external links in their search for basic information. The simplicity of this explanation has an appeal, yet it has to be tempered by WPM readers’ tendency to hover over links and click on footnotes, suggesting some level of time and interest in the sources of evidence, while a comparison with W readers points to their much greater frequency on mobile devices of clicking on external links where time might also be assumed to be an issue, with more on this is #4 below.
2. Readers may be deterred from clicking on external WPM links given that more than a quarter of the links lead to research works, of which only a fifth are publicly accessible, with earlier studies finding that encountering a paywall discourages further inquiry (Moorhead et al., 2016; Maggio et al., 2016). This suggests that something approaching universal open access to all of the research literature (along with the promotion of this fact) may be necessary, if not sufficient, to ensure that WPM readers, including those in the health professions, will pursue the greater educational advantage in clicking the external links found on WPM pages.
3. Readers may lack the background to read biomedical literature, while health professionals may have little experience in working with the research reviews favored by WPM editors, as well as other studies (although, as noted, some medical education programs include training in Wikipedia editing). If everyone is a potential or even likely user of WPM at some point, then educating people at various points in their schooling on how Wikipedia is constructed and on the roles that references play, would be helpful.
4. Readers may find the multiple links used with each research reference and other aspects of the mobile interface, including the absence of the open access icon, less encouraging than readers of the rest of W. Improvements on this front go hand in hand with continuing growth of open access to the research cited in WPM.

## Limitations

Among the limitations of this study is the way in which the count for “hovering” (defined as a reader’s cursor lingering over a link for 1,000 milliseconds or more, revealing a rollover label) captures both intentional and unintentional acts. So while the reported number of hovers is of limited value, there is no reason to believe that the incidental hovers would differ between WPM and W readers. As well, with the page view data, various strategies were used to exclude pages visited by bots, but the limited effectiveness of these strategies is a known weakness in Wikipedia’s infrastructure. It, again, suggests a tempering of the page view counts but not the ratios between WPM and W in this regard. This is similarly the case, with the exclusion from this study of interlinks that reduced external link counts, as described in the methods. Lastly, this study focuses only on the English-languge version of Wikipedia and therefore our findings are limited to this version of the encyclopedia.

## Conclusion

Through the cooperation and support of the Wikimedia Foundation, this study has compared reader engagement with external links over a 32-day period for those examining pages maintained by WikiProject Medicine and those reading others parts of Wikipedia. The distinctive qualities of WPM pages, and of reader behaviors with those pages during the period of study speak to a much stronger authorization role played by citations used in building WPM pages than for the citations serving as a gateway for readers to learn further from the works cited.

Further studies along this line would do well to seek a better understanding of whether readers find that viewing the citation itself provides sufficient “evidence” that this is an authority to be tested, or whether the potential for further learning could be facilitated through a number of design experiments involving the points raised above, possibly including (a) providing complete open access, (b) training in learning from cited works, and (c) mobile interfaces for calling up and reading research studies. Additional studies could be conducted on the access question by working with readers who have access to sources through university affiliations and with those who do not, the nature of the sources that readers seek out, from reviews of basic research to clinical trials to forms of patient-centered outcomes research, and the type of Wikipedia content associated with greater exploration of sources.

## Acknowledgements

The authors wish to acknowledge the value of this opportunity to collaborate with the WMF research team, which began in 2018 with the posting of a <u>research proposal</u> to Wikimedia’s Meta-Wiki. Members of the community, including members of WPM, were invited to provide feedback, supported by teleconferences and in-person meetings with the WMF research team on infrastructure, parameters of data use (e.g., best practices for accessing the data securely; the duration of data access), and the expectations of the Wikipedia community. As well, this collaboration extended to members of the Data Science Lab at École polytechnique fédérale de Lausanne (EPFL) who designed the CitationPageload schema, and with whom the authors developed the CitationUsage schema.

